# OOCYTE AND EMBRYO CULTURE UNDER OIL PROFOUNDLY ALTERS EFFECTIVE CONCENTRATIONS OF SMALL MOLECULE INHIBITORS

**DOI:** 10.1101/2023.11.10.566607

**Authors:** Gaudeline Rémillard Labrosse, Sydney Cohen, Éliane Boucher, Kéryanne Gagnon, Filip Vasilev, Aleksandar I Mihajlović, Greg FitzHarris

## Abstract

Culture of oocytes and embryos in media under oil is a cornerstone of fertility treatment, and extensively employed in experimental investigation of early mammalian development. It has been noted anecdotally by some that certain small molecule inhibitors might lose activity in oil-covered culture systems, presumably by drug partitioning into the oil. Here we took a pseudo-pharmacological approach to appraise this formally. Using different culture dish designs with defined media:oil volume ratios, we show that the EC_50_ of the widely employed microtubule poison nocodazole shifts as a function of the media:oil ratio, such that nocodazole concentrations that prevent cell division in oil-free culture fail to in oil-covered media drops. Relatively subtle changes in culture dish design lead to measurable changes in EC_50_. This effect is not specific to one type of culture oil, and can be readily observed both in oocyte and embryo culture experiments. We subsequently applied a similar approach to a small panel of widely employed cell cycle-related inhibitors, finding that most lose activity in standard oil-covered oocyte/embryo culture systems. Our data suggest that loss of small molecule activity in oil-covered oocyte and embryo culture is a widespread phenomenon with potentially far-reaching implications for data reproducibility, and we recommend avoiding oil-covered culture for experiments employing inhibitors/drugs wherever possible.

## INTRODUCTION

Oocyte and embryo culture in fertility clinics and research labs is routinely performed in plastic petri dishes in drops of media under a covering layer of oil to prevent contamination and evaporation, often termed the ‘closed system’^1–3^. Culture under oil allows dishes to be moved easily within the lab, permits longer-term manipulation out of the incubator such as for micromanipulation or imaging of live oocytes/embryos, and enables ex-vivo experimental access that has been fundamental in understanding early development. Typical dish setups in many labs involve multiple ∼20-50μl drops of culture media under commercially-available mineral oils in plastic petri dishes of diameter 3-5 cm, but many variations on this standard setup are used in different experimental contexts (discussed below). Extensive investigation and development takes place to ensure oils are embryo culture-safe, such as examination of the rate of media evaporation using different oils, and reducing the extent to which oils introduce pathogens or contaminants^4–6^. Here we document a further hazard of culture oil with important consequences in oocyte and embryo studies.

The past few decades have seen major advances in our understanding of oogenesis and early embryogenesis owing to the ability to pair ex-vivo culture with specific experimental interventions such as conditional transgenics^7,8^, acute gene knockdown approaches^9,10^, and use of small molecule inhibitors (‘drugs’) to interrogate molecular mechanisms^11,12^. In this context it has occasionally been noted by some investigators that they avoided oil-covered culture when using some drugs that appear to lose activity under oil, presumably a result of the ability of the drug to partition into the oil. However, a formal appraisal of the effect of oil on drug activities has not to our knowledge been presented, and whether the phenomenon is limited to just a few niche drugs, or impacts diverse and commonly used inhibitors, is unclear.

Here we quantitatively assessed the impact of oil upon the effectiveness of several cell-cycle-related inhibitors that are widely used in oocyte and embryo research. Our results show that changing media:oil ratio profoundly impacts effective drug concentrations of many, and suggest that this is a widespread phenomenon. We argue that the impacts of this upon interpretation of some published data may be significant, and make proposals for experimental design and data reporting in such experiments.

## RESULTS

### Media:oil ratios change the EC_50_ of nocodazole in oocyte maturation experiments

We reasoned that if oil reduces the effectiveness of inhibitors, then this should be quantitively demonstrable by altering the media:oil ratio in the culture dish. In this study we used a standardized series of dish setups, as illustrated in Fig1, all employing the same 35mm diameter plastic dish. At one extreme, we employed an oil-free culture dish wherein oocytes were cultured in 2ml of media in a humidified incubator in the complete absence of oil. Our groups also included one 20μl drop under 2ml oil, and ten 20μl drops under 2ml oil (ie 1:100 and 1:10 media:oil ratio respectively), reminiscent of many standard culture systems. At the other extreme we included a setup in which 2μl of media were placed under 2ml of oil (1:1000 media:oil ratio). High media:oil ratios such as this are often employed for live imaging experiments where it is important for oocytes/embryos to remain immobile, co-culture experiments, and also arise in micromanipulation studies where small media drops are used in oversized oil-filled dishes. Throughout the study we focused on well characterized inhibitors that are documented to prevent first polar body extrusion at the end of oocyte maturation. We first centered on nocodazole, a mitotic/meiotic spindle poison that causes spindle disassembly by buffering free tubulin, and thus at high concentrations prevents polar body extrusion by activating the Spindle Checkpoint^13,14^. Nocodazole is widely employed to explore cell cycle regulation in oocytes and embryos, and used routinely in micromanipulation studies to soften the cytoplasm for enucleation^15,16^. Importantly, to our knowledge, nocodazole has not previously been noted to lose activity under oil.

Nocodazole is well established to prevent first polar body extrusion during oocyte maturation by depolymerizing the spindle and therefore activating the spindle assembly checkpoint^9,14,17^. To establish an accurate EC_50_ for this effect, we initially cultured oocytes in a range of nocodazole concentrations in the absence of oil. We found that the EC_50_ for prevention of polar body extrusion in the complete absence of oil was 50.7nM, 100nM nocodazole enforcing a 100% block to Pb1 extrusion. However, EC_50_ was substantially shifted in oil-covered culture (Fig1). Notably, in the 1:100 media:oil dish setup, which reflects standard culture, the EC_50_ was shifted to 88.4nM, with 50% of oocytes extruding Pb1. Strikingly, in the 1:1000 dish setup analogous to that used in many labs for live imaging, all oocytes extruded Pb1 even at 100nM nocodazole (EC_50_ of 785nM). Thus, the EC_50_ of nocodazole for preventing Pb1 extrusion shifted more than tenfold across a range of media:oil ratios reflective of commonly used experimental setups, such that concentrations of nocodazole that prevent polar body extrusion in the absence of oil fail to do so in certain oil-covered dish setups.

Many different embryo culture oils are commercially available and used by various investigators. Therefore, to determine whether this effect was specific to one type of embryo culture oil, we cultured oocytes in 100nM nocodazole at different media:oil ratios using two different commercially available oils, one heavy oil and one light oil. Both oils are compatible with complete preimplantation embryo development in our lab (not shown). Notably, polar body extrusion occurred at higher media:oil ratios despite the presence of nocodazole, regardless of oil type (Fig2). Thus, media:oil ratios used commonly in standard experimental setups substantially impact the EC_50_ of nocodazole for preventing first polar body extrusion, and this is not specific to one type of oil.

### Media:oil ratio alters the EC_50_ of nocodazole during embryo development

To determine whether the same phenomenon could be observed in preimplantation embryos, we collected 2-cell embryos from mated females, and cultured them in the presence of nocodazole in similar dish setups as described above. Analogous to Pb1 extrusion, in the absence of oil, progression to the 4-cell stage was completely inhibited by 100nM nocodazole, embryos instead arresting in M-phase of the 2-4 cell division, consistent with the activation of the spindle assembly checkpoint as expected (Fig3). Notably however, some cells were able to divide in 100nM nocodazole in a single 20μl drop under oil (1:100), and only 6% of cells remained M-phase arrested in 2μl drops under oil (1:1000). Most cells either divided to make normal 4 cell stage cells, or underwent chaotic cell divisions, suggesting low level spindle disruption that failed to activate SAC. Thus the loss of activity under oil of nocodazole the can prevent activation of the SAC in oocytes can also be observed in embryo culture experiments.

### Impact of oil on diverse cell cycle inhibitors

Our pseudo-pharmacological analysis of the impact of media:oil ratios across a concentration range of nocodazole clearly demonstrates the impact of oil on the EC_50_ for preventing the completion of cell division, either in meiosis or mitosis. However, assembling the data presented in Fig1D alone required 1367 mouse oocytes across dozens of experimental days. We therefore sought a simplified approach that could feasibly be used by labs to test the effect of oil on a given drug. We decided to first establish the concentration-dependency for Pb1 extrusion of a given drug in the complete absence of oil, and then use a minimum effective concentration to investigate the impact of different media:oil ratios. Although this approach does not allow formal appraisal of EC_50_ shift, it provides a quantifiable indication of the extent to which drug activity is lost under oil (Fig4). We applied this approach to an additional five drugs that have been heavily employed in mouse oocyte studies:-the proteasome inhibitor MG132^18^, the kinesin-5 inhibitors STLC and Monastrol^7^, the APC inhibitor APCin^19^, and the CDK1 inhibitor Roscovitine. In all cases we added the drugs at the time of IBMX washout, as for nocodazole. APCin, MG132, STLC and Monastrol are all compatible with GVBD but prevent Pb1 extrusion, similar to nocodazole. Roscovitine prevents GVBD, and thus our analyses were on the ability of oil to permit GVBD in the presence of roscovitine. Strikingly, as for nocodazole, the ability to prevent Pb1 extrusion was substantially reduced for MG132, STLC and monastrol. Oil potently prevented roscovitine from inhibiting GVBD, even the 1:4 media:oil dish setup permitting GVBD in ∼50% of cases. Thus, oil coverings potently inactivate a range of cell cycle related drugs.

In contrast, it was noteworthy that 50μM APCin retained the ability to prevent Pb1 extrusion even at high media:oil ratios. We wondered whether this indicated a slower loss of activity, and perhaps loss of activity might become evident over a longer time-course. We therefore examined APCin after a 24h pre-incubation under oil, and found that the ability to prevent Pb1 extrusion was preserved. Thus, although nocodazole, MG132, STLC, monastrol, and roscovitine clearly lose activity under oil, we were unable to find evidence that APCin activity is lost, even after long-term media-oil contact.

Finally, we wondered whether drugs that lose activity under oil may do so even more profoundly after a 24h incubation. To test this we examined the impact of a 24h preincubation under oil upon MG132, one of the drugs that most profoundly lost activity even in our standard experimental setup (2h preincubation under oil). We found that MG132 was even further inactivated by a 24h incubation under oil, such that ∼60% of oocytes extruded Pb1 in the 1:100 dish setup compared to only ∼10% in the standard 2h experiment (all at 5μM MG132). This indicates that even the timing of dish setup (eg, whether dishes are prepared the night before) affects drug EC_50_s.

To summarise, our analysis of a panel of 6 drugs suggests that although some specific drugs may retain their activity under oil, as exemplified by APCin, five of the six we tested are very clearly inactivated by a covering layer of oil, and that the extent of this inactivation can be dependent upon time spent under oil.

## DISCUSSION

Here we have used a conceptually simple experimental approach to demonstrate that the effective concentration of an array of commonly used cell cycle drugs is dramatically changed by culturing under oil. Although we have not formally measured the presence of drugs within oil after culture, the simplest interpretation is that each drug partitions into the oil, and the oil acts as a sink. Some level of hydrophobicity is necessary for most drugs to enter cells, and *LogP* values that provide a broad indication of oil solubility based on water/octanol partitioning and are provided with most reagents would tend to support the notion that these drugs would partition into oil.

In some cases partial drug loss under oil may have little impact on broad experimental conclusions, particularly where supra-maximal concentrations are used to elicit well characterised effects. However, there are several types of conclusions that are more precarious. For example, ‘negative results’ in which a drug appears to have no impact upon a cellular process, or circumstances in which unexpectedly high concentrations are needed to elicit expected effects, may warrant revisiting. Moreover, interpretations of which molecular species are being inhibited by selective inhibitors at specific concentrations should be interpreted with extreme caution if oil-covered culture was employed. Instances in which different inhibitor concentrations were required between studies to achieve a given phenotype could be explained by different dish setups and even the timing of their preparation.

Although here we have formally examined the loss of activity under oil of only 6 drugs, there are strong clues that many others behave similarly. Blegini and Schindler^20^ have noted that they avoid oil for Aurora-Kinase inhibitors. The Wassmann group noted that when using the MPS1/AurK inhibitor Reversine they supplemented the culture oil with drug^21^. Halet et al used the PI3K inhibitor LY294002 in oil free conditions, noting a dramatic reduction in the required concentration to prevent preimplantation embryo development compared to other studies^22,23^. Doubtless many other examples exist. Other drugs that we have anecdotally observed in our lab to lose activity under oil include the CENPE inhibitor GSK923295^24^, the APC^cdh1^ inhibitor Protame^24^, and the myosin ATPase blocker Blebbistatin^25^. Thus while we believe the present study is the first to formally quantitate and highlight the effect, loss of drug activity under oil has certainly been noted by others, and is likely a widespread phenomenon.

To conclude, we advise caution when employing small molecule inhibitors, avoiding oil-covered culture wherever possible. Where the use of oil is unavoidable given the experimental context, in some live imaging studies for example, thorough testing should be carried out to demonstrate that the media:oil ratio employed is far below the threshold where results are affected. Most importantly, detailed experimental information including exact media:oil ratios and timing of dish setup should be clearly reported when drugs are used under oil. As increasing importance is rightly placed upon data reproducibility, elimination of factors that inadvertently change experimental conditions in a manner that could critically alter results is paramount.

## MATERIALS AND METHODS

### Oocyte and embryo collection

Mouse oocytes were collected at the GV stage from the ovaries of CD1 females (Charles River Crl:CD1(ICR) 022) aged 6-12 weeks, after intraperitoneal injection of 5 IU pregnant mare serum gonadotropin (PMSG; Aviva system biotech OPPA01037). Oocytes were collected in M2 media containing 200 μM 3-isobutyl-I-methylxanthine (IBMX; Sigma I5879). Following collection oocytes were kept in M16 medium supplemented with IBMX to prevent GVBD (Wisent 311-630-QL) 37°C under 5% CO_2_, prior to transfer to experimental dishes (see below). Mouse two-cell embryos were collected from the oviducts of CD1 females aged 6-12 weeks, following intraperitoneal injection of 5 IU pregnant mare serum gonadotropin (∼ 90 hours pre-collect) and 5 IU human chorionic gonadotropin (hCG; Sigma; ∼40-48 hours pre-collect) and mating with BDF1 male mice (Jackson 100006). Embryos were collected in M2 media and cultured in KSOM (Wisent 003-026-XL) under 5% CO_2_ at 37 °C. All animal experiments were performed in accordance with relevant CIPA (Comité institutionnel de protection des animaux - CHUM) guidelines and regulations under protocol IP22054GFs.

### Small molecule treatments

Following collection oocytes were incubated in M16 + IBMX for 1-3 hours to allow zona release^26^ and a final selection of healthy fully-grown oocytes was performed. To test the effect of oil on inhibitors, 35-mm plastic culture dishes (Sarstedt 83900) were prepared with different media:oil ratios as indicated in Figure 1 (‘experimental dishes’). Importantly, experimental dishes were prepared exactly 2 hours prior to the addition of oocytes/embryos for examination of the effect of drugs/oil in all cases, with the exception of Figures 4D and E which also included a group in which the dishes were assembled 24 hours prior. Light oil (Sigma M8410) was used for all experiments, and heavy oil (Sigma 330760) was used in addition in Figure 2. The media:oil ratios used in experimental dishes were: no oil (2mL media without oil); 1:4 (1 × 500 uL media covered with 2 mL oil); 1:10 (10 × 20 uL media covered with 2 mL oil); 1:100 (1 × 20 uL media covered with 2 mL oil) and 1:1000 (1 × 2 uL media covered with 2 mL oil)). For embryo experiments, two-cell embryos were collected and then transferred in inhibitors 2 hours after collection, which was again exactly 2h after dish setup.

**FIGURE 1.**
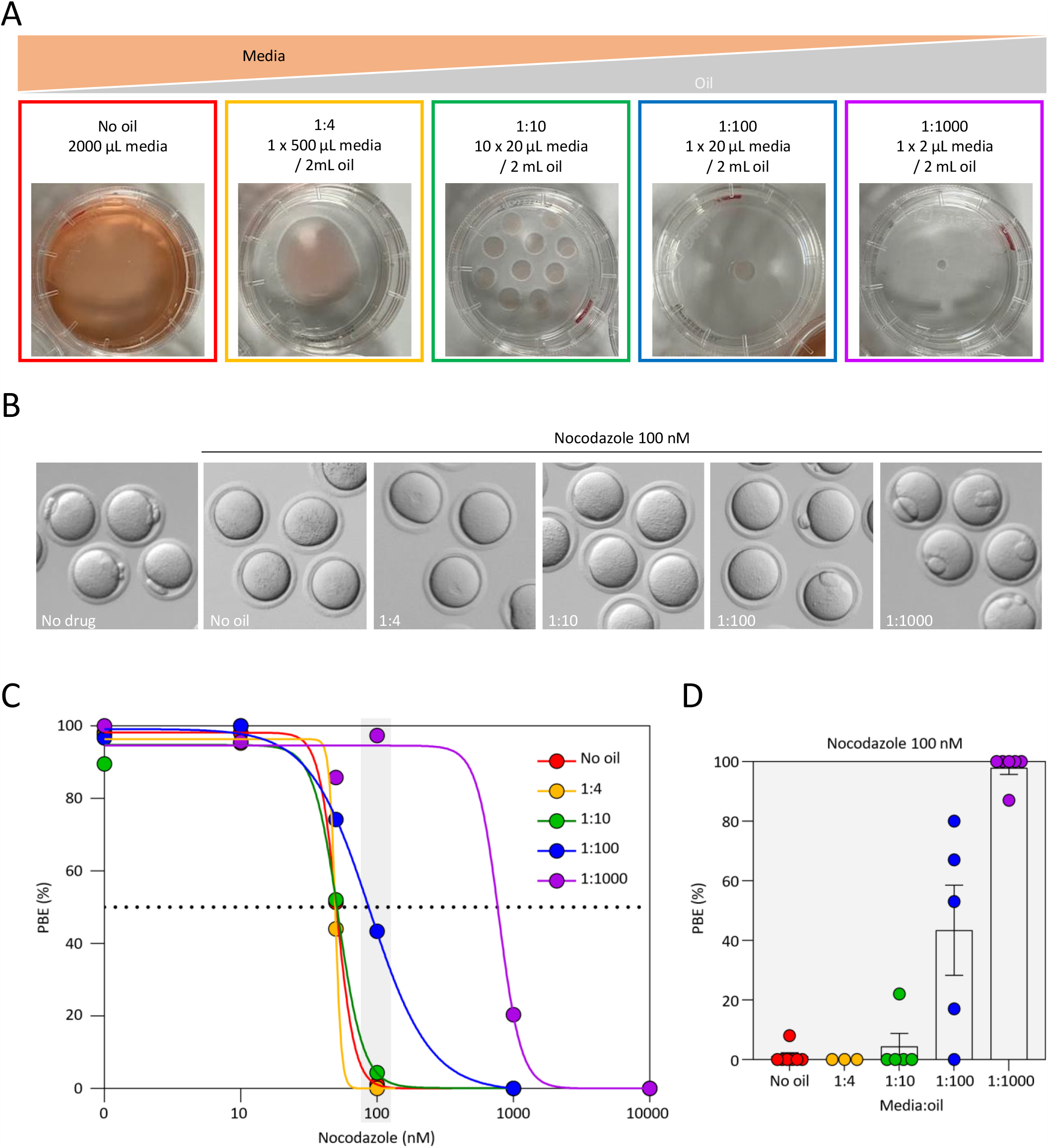
Effect of media:oil ratio on the EC_50_ of nocodazole during oocyte maturation. **(A)** Schematic representation of the experimental design to address impact of media:oil ratio on drug effects. Conditions were: no oil (2mL media without oil); 1:4 (1 × 500 uL media covered with 2 mL oil); 1:10 (10 × 20 uL media covered with 2 mL oil); 1:100 (1 × 20 uL media covered with 2 mL oil) and 1:1000 (1 × 2 uL media covered with 2 mL oil). **(B)** Bright-field images illustrating different outcomes of PBE in oocytes incubated in 100 nM nocodazole for different media:oil ratios. **(C)** Dose-response curves for oocytes incubated in varying concentrations of nocodazole in different media:oil conditions. For this graph, each data point is an average of 3-8 replicates, ∼10 oocytes per replicate, n=1367 oocytes in total. **(D)** Bar chart representation highlighting impact of different media:oil ratios for oocytes incubated in 100nM nocodazole. Data from same experiments as Fig1D; each data point is a replicate of ∼10 oocytes, n=332 oocytes in total. Bars represent mean ± SEM.

**FIGURE 2.**
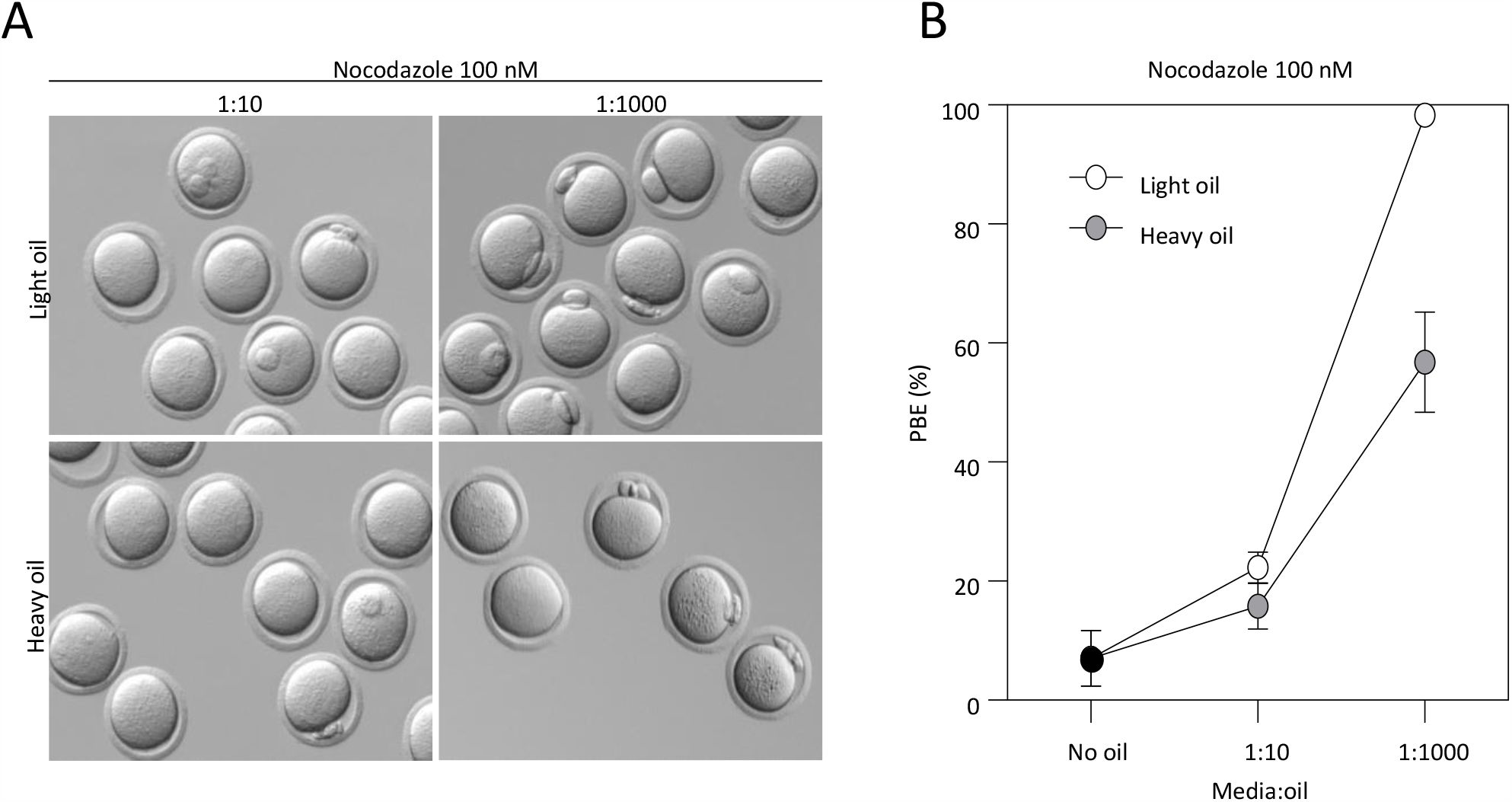
Loss of nocodazole activity is not specific to only one type of oil. **(A)** Bright-field images illustrating outcomes for oocytes incubated in 100nM nocodazole for different media:oil ratios in light versus heavy mineral oil. **(B)** Graph representing changing PBE rate for oocytes incubated in 100nM nocodazole in different media:oil conditions comparing two oil types (light vs heavy mineral oil). For this graph, each data point is an average of 4 replicates, ∼10 oocytes per replicate, n=223 oocytes in total. Bars represent mean ± SEM.

**FIGURE 3.**
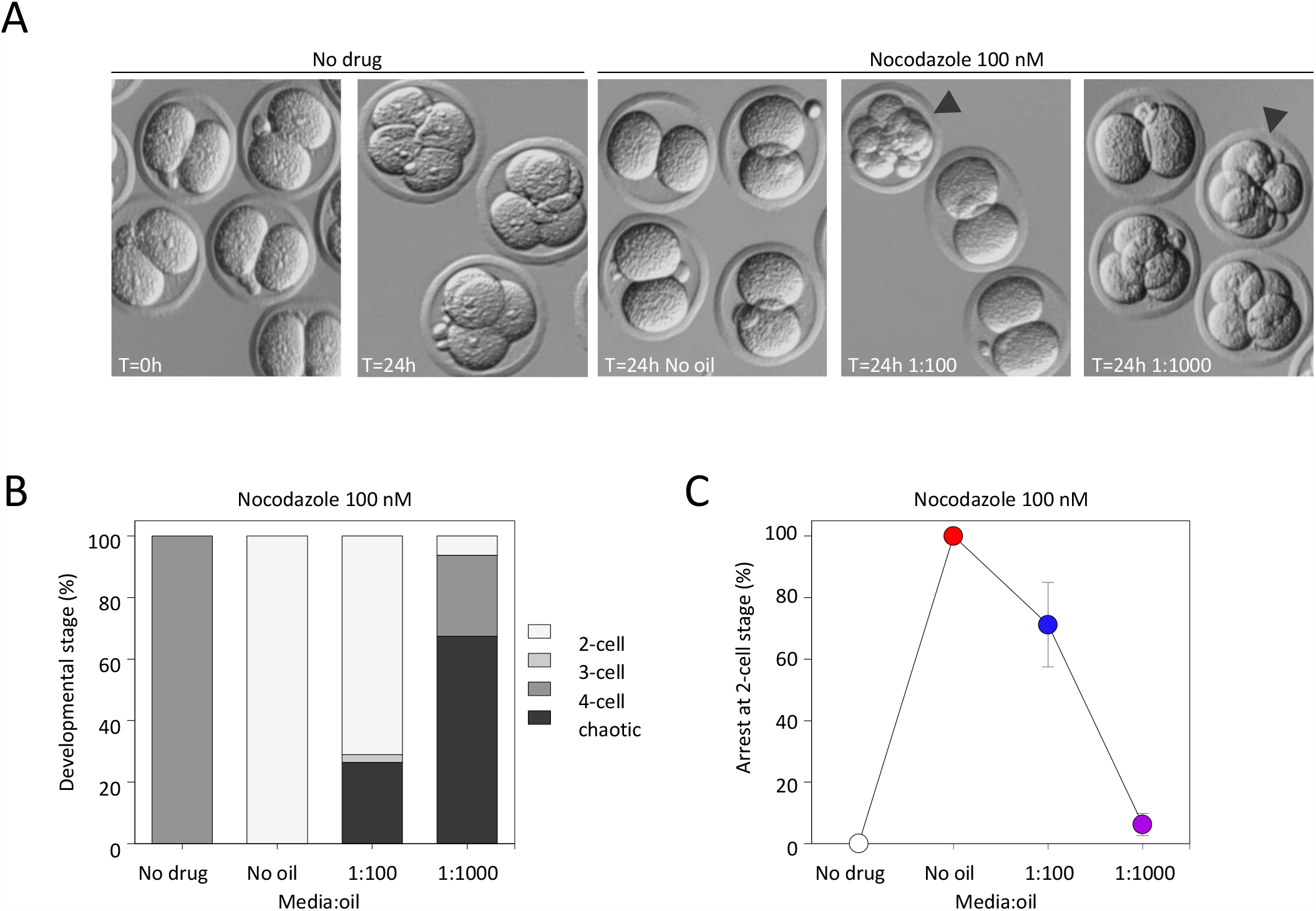
Impact of oil on the EC_50_ of nocodazole during embryo development. **(A)** Bright-field representative images illustrating appearance of embryos incubated in 100nM nocodazole with different media:oil ratio conditions. Arrowheads indicate embryos characterized as chaotic. **(B)** For each media:oil condition embryos were categorized in sub groups depending on the blastomere count (2-cell, 3-cell, 4-cell and chaotic). **(C)** Arrest at 2-cell stage was measured after embryos were incubated in 100 nM nocodazole in different media:oil ratios. Bars represent mean ±SEM. For B and C, each data point is an average mean of 4 replicates, 120 embryos in total.

**FIGURE 4.**
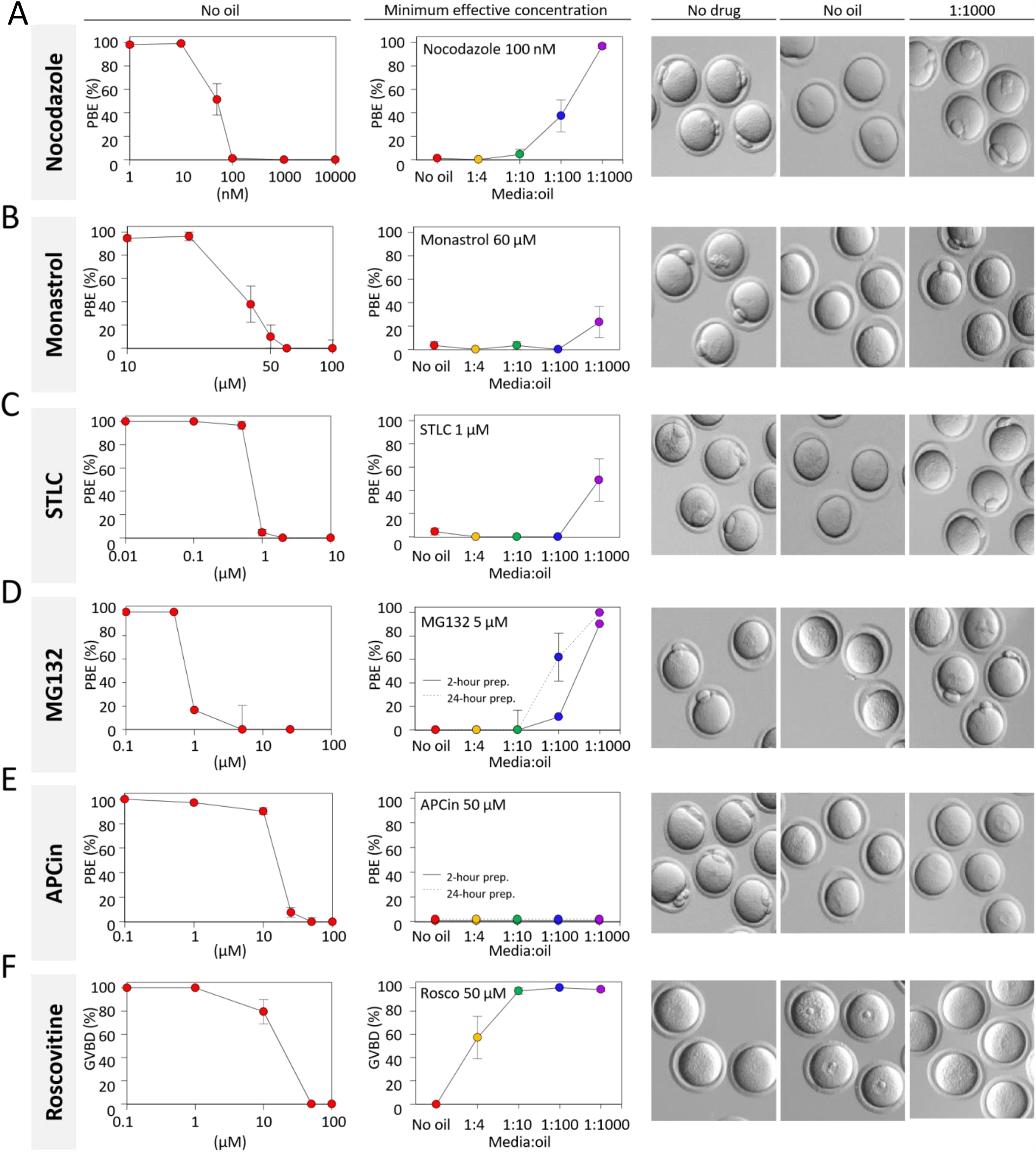
Oil covered culture affects the action of many inhibitors. Polar body extrusion (PBE) was assessed following incubation of GV oocytes with **(A)** nocodazole; **(B)** monastrol; **(C)** STLC; **(D)** MG132; **(E)** APCin. Germinal vesicle breakdown (GVBD) was evaluated with GV oocytes treated with **(F)** roscovitine. Oocytes were first incubated in varying concentrations of inhibitors without oil in order to determine the effective concentration of these drugs (ie, concentration-response curves). In the central column, the lowest maximally effective concentration was used to analyze the impact of media:oil ratios. Dishes with media containing inhibitors ± oil were prepared and pre-equilibrated at 37°C with 5% CO2 for 2 hours prior to oocytes being transfer to them, as elsewhere in the study, and for MG132 (D) and APCin (E), a parallel set of dishes were additionally pre-equilibrated for 24 hours prior to oocytes incubation. On the right, bright-field images represent oocytes status when evaluating PBE and GVBD. Each data point is the mean of a minimum of 3 replicates, ∼10 oocytes/replicate, a total of 2245 oocytes throughout the figure. Bars represent mean ± SEM.

Oocyte maturation was triggered by washing oocytes into IBMX-free media. Immediately thereafter, oocytes were transferred to into the experimental dishes. Oocytes or embryos was washed through 4 drops of media with inhibitor in a similar setup (concentration, and media:oil ratio) as the final experimental dish, and then transferred to the experimental dish. On any given experimental day there was an additional control group of oocytes or embryos incubated entirely without inhibitors - data was included in the final dataset from any experimental day only if >90% development (GVBD, PBE, or 2-4 cell division) was observed. Inhibitors used in this project were: nocodazole (Calbiochem/Millipore 487928); monastrol (Calbiochem/Millipore 475879); S-Trityl-L-cysteine (STLC; Tocris 2191); MG132 (Calbiochem/Millipore 474790); APCin (Tocris 5747) and roscovitine (Sigma R7772). Effects of the treatment was assessed 16-18 hours post transfer into inhibitors. Bright-field images were captured using a Leica M165C dissection scope equipped with a camera (Camera Opti-Vision 4K LITE-8MP Opti-Tech Scientific).

## ACKNOWLEDGEMENTS

We thank François Proulx, Karen Schindler, and Guillaume Halet for valuable discussions and for critical reading of the manuscript. This research was supported by the grants from Canadian Institute for Health Research (CIHR), Foundation Jean-Louis Levesque, and Natural Sciences and Engineering Research Council of Canada (NSERC) to GF. EB received a scholarship from McGill University’s Centre for Research in Reproduction and Development (CRRD), and SC received a scholarship from Université de Montréal Centre de recherche en reproduction et fertilité (CRRF).

## AUTHOR CONTRIBUTIONS

Gaudeline Rémillard Labrosse, Sydney Cohen, Éliane Boucher, Kéryanne Gagnon: Equal contributions in data curation; formal analysis; validation; investigation; writing review and editing manuscript.

Filip Vasilev and Aleksandar I Mihajlović: data curation; formal analysis; validation; investigation; visualization; writing reviewing and editing.

Greg FitzHarris: conceptualization; supervision; funding acquisition; data curation; formal analysis; validation; writing–original draft; writing–review and editing.

